# Bilingualism Protects Domain-Specific Cognitive Function in Mandarin-Speaking Older Adults

**DOI:** 10.64898/2026.02.02.703396

**Authors:** Yulania Wang, Oceanna Li, Feng Vankee Lin, Meishan Ai

## Abstract

**Objectives:** This study examined whether bilingualism is associated with episodic memory (EM) and executive function (EF) in older Mandarin-speaking adults and whether associations differ by clinical diagnosis.

**Methods:** 189 Mandarin-speaking older adults completed Mandarin-administered neuropsychological testing, brain MRI, and clinical diagnosis. English proficiency test was administered to determine whether they were bilinguals or monolinguals. Sensitivity analyses were conducted in an education-matched subgroup (16 years).

**Results:** Bilingualism was associated with higher EF, with a significant bilingualism × diagnosis interaction indicating larger bilingual advantages among cognitively impaired participants. However, bilingualism was not associated with EM. Findings were preserved in education-matched analysis.

**Conclusions:** Results support a domain-specific association between bilingualism and executive function in later life, consistent with cognitive maintenance mechanisms preferentially supporting executive processes rather than global protection.

## INTRODUCTION

Bilingual language experience has been shown to be protective for cognitive function in late life and has been proposed as a cognitive reserve factor that buffers against neuropathology.^1^ Managing two languages across the lifespan places sustained demands on control and selection processes supported by domain-general executive function. Additionally, neurocognitive evidence indicates that language control selectively engages and enhances brain regions associated with executive function.^2^ Consistently, several studies reported delayed clinical manifestation of dementia symptoms among bilingual individuals, suggesting preserved functional performance despite underlying pathology.^3,4^ However, empirical findings remain mixed. Bilingualism-related advantages have been reported more consistently for specific executive function processes, but not for other cognitive domains, and are variably expressed across populations differing in cognitive status, language experience, and sociocultural context.^5,6^

The inconsistency is potentially caused by methodological heterogeneity, including variability in cognitive tasks, inconsistent second language of testing, and limited consideration of clinical cognitive status.^7^ Reviews highlight inconsistent assessment language, insufficient control of demographic confounding factors, and clinical heterogeneity as major obstacles in the bilingualism and aging literature.^6^ Additionally, most previous studies examine Indo-European language pairs, typically English–Spanish, which share substantial overlap in phonology, orthography, and morphosyntax. Findings from these populations may therefore not fully generalize to bilingual experiences involving more typologically distinct languages. For example, Mandarin–English bilingualism might require greater working memory capacity to manage and switch between logographic and alphabetic systems,^8^ potentially exercising different cognitive processes. Studying bilingualism in this context offers unique insights into how bilingualism of distinct language systems enhances cognitive function.

We aim to addresses these gaps by examining the association between bilingualism and cognitive function in older adults who speak Mandarin primarily and English secondarily. Specifically, this study primarily focused on Episodic Memory (EM) and Executive Function (EF) in older Mandarin-speaking adults across cognitive diagnosis status, as the two domains are most susceptible to age-related decline and neuropathology, but with partially dissociable neural substrates and trajectories.^9,10^ Moreover, as a key confounding variable, education attainment was treated as a covariate and education-matched sensitivity test was conducted in the examination of bilingual language experience. Together, these analyses clarify how bilingual experience relates to cognitive performance across domains and cognitive diagnostic status, contributing to understanding how life experience influences cognitive aging.

## METHODS

### Study cohort (COAST)

The Chinese Older Adult Study of Cognition and Technology (COAST) is an ongoing community-based study designed to investigate cognitive aging among Mandarin-speaking older adults, with particular emphasis on bilingual Mandarin–English speakers. COAST addresses longstanding challenges in cognitive assessment for Chinese-speaking populations by administering culturally and linguistically adapted neuropsychological measures in Mandarin. 189 older adults were recruited from the New York City/New Jersey (NYC/NJ) and San Francisco Bay (SF Bay) areas between 2023 and 2025 through community outreach efforts, including flyers, community events, Chinese-language newspapers, WeChat, and word of mouth. Eligible participants were aged 50 years or older, fluent in Mandarin, and self-identified as ethnically Chinese or Taiwanese. All participants underwent comprehensive neuropsychological assessment administered in Mandarin, structural brain MRI, clinical evaluation, and blood collection. Ethical approval was obtained from the Rutgers Biomedical and Health Sciences and Stanford University Institutional Review Boards. All participants provided written informed consent in their preferred language (simplified Chinese, traditional Chinese, or English). Capacity to consent was assessed using a modified University of California San Diego Brief Assessment of Capacity to Consent, with scores above 14 required for enrollment.

Demographic, socioeconomic, and immigration-related information, including age, sex, education, was collected via structured interview. Medical history and medication use were documented. For most participants, a knowledgeable study partner completed the Clinical Dementia Rating (CDR) interview. The current analyses included 189 participants who completed neuropsychological testing and consensus diagnosis as of September 11, 2025.

### Bilingualism

Language proficiency was assessed using brief, standardized web-based tests. English proficiency was assessed via two independent tests: a reading and writing test (Transparent Language, Nashua, NH, USA) and a reading and listening test (EF SET, Signum International AG, Zurich, Switzerland). The final English proficiency score was coded according to the Common European Framework of Reference for Languages (CEFR) as Basic (A1/A2), Independent (B1/B2), or Proficient (C1/C2). Participants were classified as bilingual if they achieved a CEFR level of B1 or higher on at least one English proficiency test. Participants below this threshold were classified as monolingual Mandarin speakers. This operationalization was selected to capture functional bilingual ability relevant to daily language use and executive language control.

### Cognitive assessments

EM and EF composite scores were obtained via a factor analysis across multiple cognitive tasks.^11^ The EM factor scores was summarized across the Craft Story task, a narrative memory task included in the Uniform Data Set, and the immediate recall (IR) and delay recall (DR) tasks. Compared with the English version, the original Chinese translation of the Craft Story contained a greater number of lexemes and longer multi-character words, resulting in longer reading time and potential interpretive ambiguity. Based on pilot testing, 12 lexemes were simplified, truncated, or substituted to reduce linguistic complexity while preserving narrative structure and memory demands. The adapted version consisted of 68 characters, comparable to the syllabic length of the English story. The IR and DR tests were scored using standardized verbatim and paraphrase criteria. All subtest scores were standardized and averaged to create a composite EM z-score, with higher values indicating better performance.

The EF factor score was summarized across Trail Making Test Part A (TMT-A), Trail Making Test Part B (TMT-B), and the Symbol Digit Substitution Test (SDST). TMT-A primarily measures processing speed and visual scanning, whereas TMT-B additionally requires alternating set-switching and cognitive control. The SDST assesses speeded attention and working memory. Raw scores for the three tests were transformed such that higher values reflected better performance, standardized, and averaged to generate a composite EF z-score. For more information about the tests and factor analysis, please refer to Hu et al.^11^

### Diagnostic formulation and neurodegeneration

Cognitive status was determined through a multidisciplinary consensus process conducted by study clinicians and investigators, following National Institute on Aging–Alzheimer’s Association (NIA-AA) criteria. Diagnoses of normal cognition (NC) or cognitive impairment (mild cognitive impairment or dementia) were assigned based on demographic and clinical history, medical information, Clinical Diagnostic Rating (global score and sum of boxes), structural MRI findings, and English-language neuropsychological data when available, consistent with Uniform Data Set 3.0 procedures. For monolingual participants without English testing, diagnosis was determined using the same consensus process incorporating Mandarin test performance and MRI findings.

AD-related neurodegeneration was indexed using AD-signature cortical thickness (ADSCT) derived from structural MRI. For information about data acquisition and preprocessing please refer to supplementary materials. Cortical surface reconstruction was performed by FreeSurfer. Regional thickness values were extracted based on the Desikan–Killiany–Tourville atlas and averaged across bilateral entorhinal cortex, fusiform gyrus, and middle and inferior temporal gyri, consistent with established AD-signature indices. Higher ADSCT values indicate greater cortical thickness and less AD-related neurodegeneration.

### Statistical Analysis

All statistical analyses were conducted in Python using the *statsmodels* and *pingouin* libraries. Analyses were performed using listwise deletion such that participants with missing data on variables required for a given model were excluded from that analysis. Multivariable linear regression model was conducted to examine the relationship between bilingualism and cognition (i.e., EF and EM), where age, sex, education, AD-signature cortical thickness, cognition diagnostic status, and study sites were controlled as covariates. Separately, interaction term between diagnostic status and bilingualism was added to the original model to examine whether bilingualism-cognition relationship differs across diagnostic groups. Model fit was evaluated using R^2^ and adjusted R^2^, and regression coefficients were estimated using ordinary least squares. To control for multiple comparisons across the two cognitive domains, p-values for bilingualism-related effects were corrected using a Bonferroni adjustment, multiplying nominal p-values by 2. Additionally, sensitivity analysis was conducted in a restricted sample with 16 years of education across both bilingual and monolingual participants, to evaluate the robustness of bilingualism effects with respect to educational differences between language groups. Within this restricted sample, the primary regression models were re-estimated using the same covariate structure except education. For descriptive visualization, we plotted EM and EF performance by diagnosis status, stratified by bilingual group. Covariate-adjusted group means with 95% confidence intervals were displayed for each bilingual group to illustrate patterns of cognitive performance across diagnostic groups. The characteristics for COAST sample was summarized in Table 1.

**Table 1.**
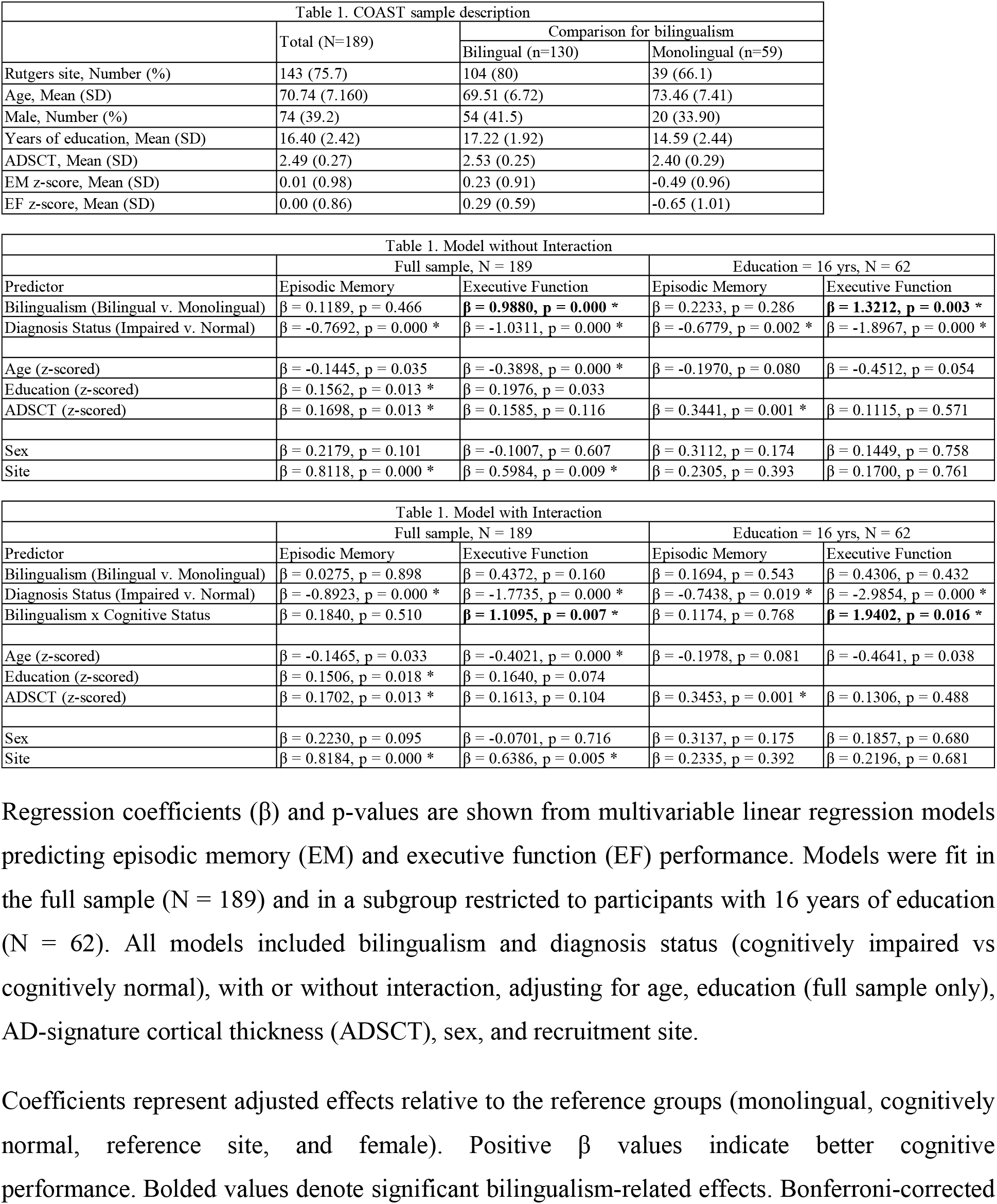
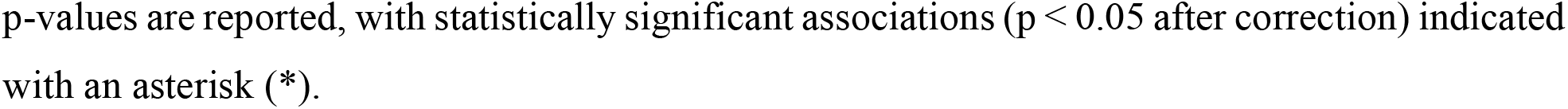
Associations of bilingualism and diagnosis status with episodic memory and executive function in the full sample and education-matched subgroup.

## RESULTS

### Main effects of bilingualism

In the full sample, bilingualism was not associated with EM performance after controlled for covariates (β = 0.1189, p = 0.466). However, bilingualism was significantly and positively associated with EF (β = 0.9880, p = 0.000; Bonferroni-corrected p < .05). The parameters of the full model were reported in Table 1.

### Interaction analyses: bilingualism × diagnostic status

To determine whether bilingualism-related differences varied by clinical context, we tested bilingualism by diagnostic status interactions for each cognitive domain.

No significant bilingualism × diagnosis status interaction was observed for EM (p = .510; Figure 1A), whereas a significant interaction was observed for EF (β = 1.11, p = .007; Bonferroni-corrected p < .05). Model estimates indicated that bilingual–monolingual differences in EF were substantially larger among cognitively impaired participants than among cognitively normal participants (Figure 1B). Specifically, cognitively impaired bilingual participants exhibited significantly better EF performance than cognitively impaired monolingual participants, whereas bilingual–monolingual differences were smaller and not statistically robust among cognitively normal individuals. The inclusion of this interaction term also improved model fit, with the adjusted R^2^ remaining high (adjusted R^2^ = .51). Additional interaction analyses showed that bilingualism did not significantly interact with age or ADSCT for EF or EM, indicating that bilingualism-related differences did not reflect altered sensitivity to aging or AD-related neurodegenerative burden.

**Figure 1.**
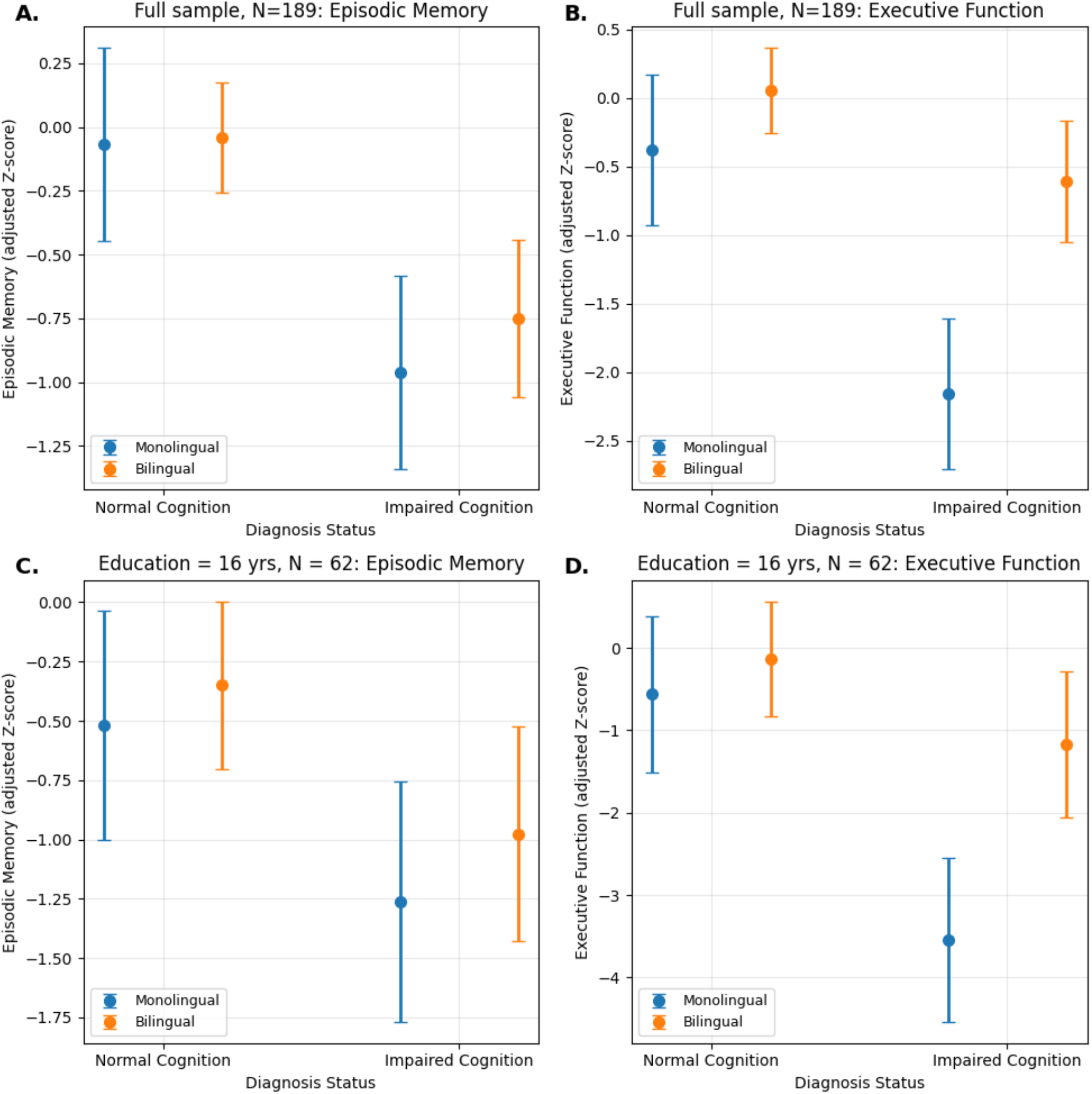
Bilingualism and diagnosis status interactions for episodic memory and executive function. Covariate-adjusted episodic memory (EM) and executive function (EF) performance as a function of bilingualism and diagnosis status. Panels show covariate-adjusted mean cognitive z-scores with 95% confidence intervals for monolingual and bilingual participants within cognitively normal and cognitively impaired diagnosis groups. Panels A and B present results for the full sample (N = 189), whereas Panels C and D show results restricted to participants with 16 years of education (N = 62). Models adjust for age, AD-signature cortical thickness, sex, and recruitment site, with education additionally included as a covariate in full-sample analyses. Monolingual and cognitively normal participants serve as reference groups in regression models.

In sensitivity analysis of education-matched sample, bilingualism × diagnostic status interaction was not significant for EM (β = 0.1174, p = 0.768, Figure 1C) but significant for EF (β = 1.94, p = 0.016). Consistently, cognitively impaired participants continued to demonstrate greater bilingual-related advantage on EF than cognitively normal participants (Figure 1D).

## DISCUSSION

This study examined whether bilingualism relates to cognitive performance in older Mandarin-speaking adults and whether such associations differ by diagnostic status after accounting for demographic factors and AD-related neurodegeneration. The findings suggest that bilingual language experience is linked more closely to EF than to EM, consistent with the view that life-experiential factors may differentially support cognitive systems in aging.

The differential associations between bilingualism and EF versus EM likely arise from the partially dissociable neural systems underlying these cognitive domains.^9,10^ Memory performance in later life is strongly constrained by neurodegenerative burden and earlier-life learning experiences.^12^ Meanwhile, EM systems depend heavily on medial temporal structures that are particularly vulnerable to Alzheimer’s disease pathology,^13^ potentially limiting the extent to which later-life experiential factors influence performance in this domain.

Moreover, this domain difference may also attribute to the cognitive processing being exercised from lifelong management of two languages. EF plays a central role in bilingual language use, as speakers must continuously select the appropriate language and switch between linguistic systems based on conversational context, potentially enhancing these processes through lifelong experience.^14^ Evidence from neuroscience study^15^ also revealed greater brain functional efficiency underlying cognitive control, a core mechanism of EF, in bilinguals comparing to monolinguals, which might support better maintained EF in later-life.

Importantly, the bilingualism-EF relationship was more apparent under conditions of cognitive impairment, further suggesting that such experiential influences may become most visible when cognitive resources are constrained. This finding was also consistent with previous reports on bilingualism and cognition.^6^ This pattern suggests the compensatory effect of bilingual experience on EF on under conditions of cognitive vulnerability.

Importantly, this study controlled the confounding effect of educational attainment by treating education as a covariate and conducting a sensitivity analysis in an education-matched sample. The consistent observation of the bilingualism-EF relationship further suggests the unique contribution of bilingual experience to EF in later-life and across cognitive diagnostic status, independent of education.

Some limitations should be considered. Future longitudinal follow-up will be valuable for understanding how bilingual language experience influences cognitive trajectories across aging. Bilingualism was summarized categorically in one dimension; a continuous multidimensional construct needs to be established in future research to precisely and comprehensively collect bilingual experience. In addition, ADSCT represents only one aspect of neurodegeneration and may not fully reflect neural processes supporting EF. Future longitudinal studies incorporating more detailed bilingual-related index and multimodal biomarkers will help clarify how bilingual experience relates to cognitive aging over time.

In summary, the present findings support a domain-specific association between bilingual language experience and EF in later life. Moreover, more apparent bilingualism-EF effect in cognitive impairment group suggests that the bilingual experience might play a compensating role in protecting EF in the presence of cognitive burden.

## Acknowledgments

We greatly appreciate the support from the National Institutes of Health. We thank all the collaborators, consultants, research scientists, and clinical coordinators from the research group of COAST study for their contributions. We especially thank all the participants for their time and efforts.

