## Supplementary Materials for "Bilingualism Protects Domain-Specific Cognitive Function in Mandarin-Speaking Older Adults"

### MRI acquisition and processing.

Structural MRI data were acquired at two sites using 3T scanners. At Rutgers University, images were collected on a Siemens Prisma scanner with a 32-channel head coil using a high-resolution 3D MP-RAGE sequence [Repetition time/Echo time (TR/TE) = 2370/2.34ms, flip angle = 8°, voxel size = 1 mm isotropic, field of view (FOV) = 256mm, 192 sagittal slices, GRAPPA acceleration factor = 2, echo spacing = 7.04ms]. At Stanford University, images were acquired on a GE SIGNA Premier scanner with a 48-channel head coil using a comparable 3D MP-RAGE sequence [Repetition time/Echo time (TR/TE) = 2320/2.94ms, flip angle = 8°, slice thickness = 1mm, FOV = 256mm, in-plane acceleration = 2, matrix = 256 × 512, resolution = 1 mm isotropic].

T1-weighted structural images from both sites were processed using the FreeSurfer recon_all pipeline, including skull-stripping, tissue segmentation, cortical surface reconstruction, and spatial normalization.
